# deeprok: highly accurate prokaryotic gene prediction through deep learning and overlap-aware decoding

**DOI:** 10.64898/2026.11.21.768321

**Authors:** Qiaohan Xu, Heng Li, Jim Shaw

## Abstract

Prokaryotic gene prediction is often considered solved, yet widely used tools still disagree on many genes, chiefly short genes and translation starts. We present deeprok, an ab initio prokaryotic gene prediction method that combines a base-level neural network with an overlap-aware decoder, and refines start-site predictions with a self-trained neural ranker. On four curated genomes distant from its training data, deeprok made 10–18% fewer exact-match errors than Prodigal and GeneMarkS-2, and on 2,925 experimentally verified starts it made 21–41% fewer start errors. Furthermore, deeprok’s base-level architecture reveals lineage-specific translation initiation signals. These results show that per-base deep learning combined with overlap-aware decoding can improve prokaryotic gene prediction, especially for resolving translation starts.

## Background

Gene prediction is the first step of genome annotation. Accurate predictions are necessary because errors propagate into downstream analysis [1]. In prokaryotes the task is often considered largely solved: genomes are densely coding, with roughly 88% of sequence annotated as coding, and genes lack introns [2,3]. Yet no single gene finder performs best across organisms, and even top-ranked tools produce conflicting gene sets [4]. A recent benchmark of annotation pipelines across more than 150,000 genomes found that no pipeline is universally optimal and called for more accurate start codon and ribosome binding site assignment [3]. In practice, two types of errors persist. The first type is false-positive gene calls, which are concentrated among short open reading frames (ORFs) because false discovery rates rise sharply as ORF length decreases [5,6]. Most gene prediction methods therefore use a minimum ORF size parameter, but this also discards genuine small proteins. The second is incorrect translation start sites: widely used tools disagree for 15–25% of genes in a genome [7]. Novel genomes often lack RNA-seq, proteomics or close homologs to check predictions against, so ab initio accuracy is critical.

Ab initio prokaryotic gene finders are built using statistical models of coding sequence. Prodigal, for example, scores ORFs using coding statistics and ribosome binding site (RBS) motifs learned from the input genome and then selects a set of ORFs by dynamic programming. Its scoring heuristics, however, were developed on a small set of well-curated genomes [8].

GeneMarkS-2 instead trains genome-specific hidden Markov models (HMMs) through unsupervised self-training. It models translation initiation by assigning each genome to one of a few predefined classes of upstream motifs, and it uses precomputed models for genes with atypical composition [9]. While both tools are popular and empirically successful, they rely on hand-crafted features and predefined start-site patterns learned from a small set of genomes.

This raises the question of whether a model that learns sequence features directly could generalize better.

Deep learning has recently emerged as a powerful approach to ab initio gene prediction in eukaryotes due to its ability to leverage large-scale databases that are now available. For example, Helixer combines convolutional and recurrent layers to predict a genic class for every base, and then derives a final set of genes in an HMM-based post-processing step [10]. Tiberius builds on this by integrating a differentiable HMM layer that is trained end-to-end with the network [11]. In both methods, the network produces per-base predictions, and the HMM then infers a single most likely gene structure. However, this approach does not transfer directly to prokaryotes, where roughly one third of genes overlap a neighboring gene [12]. Because each position has a single hidden state, the HMM cannot represent two genes that share bases.

Existing deep learning methods for prokaryotes instead focus on using neural networks to score ORFs. For example, Balrog applies a temporal convolutional network to the translated protein sequence of each candidate ORF [13]. It then selects genes through a graph search whose edge weights combine gene and start-site scores, a start-codon bonus and overlap penalties, without modeling the intergenic sequence between genes. Similarly, GeneLM fine-tunes a DNABERT genomic language model to classify candidate ORFs and then their start sites [14]. To our knowledge, no prokaryotic gene caller yet combines per-base predictions from a neural network with explicit, overlap-aware scoring of candidate genes. A neural network that outputs per-base predictions makes the model more inspectable, as one can examine which positions and signals affect gene prediction. This also allows users to examine coding probabilities of specific sites of interest when no external evidence is available for validation.

Here we present deeprok, an ab initio gene caller for prokaryotes. In deeprok, a convolutional and bidirectional LSTM network trained on 666 genomes predicts per-base labels. Candidate genes are then scored under an HMM probability model of gene structure, and a gene set is selected by dynamic programming that accommodates overlapping genes. To improve start-site assignment, we also adopt the idea of self-training. In contrast to approaches that use predefined RBS categories for prediction, deeprok uses a distinct second-pass neural network that learns start-site features from the sequence of its own predictions and then ranks putative start sites. In our benchmarks, deeprok had the highest exact-match accuracy on four curated genomes and the most accurate start calls on 2,925 experimentally verified genes, while making fewer unexplained false positives than either widely used caller.

## Results

### Overview of deeprok

deeprok is a deep learning gene finder that annotates protein-coding genes in bacterial and archaeal genomes directly from genomic DNA (Fig. 1). The architecture of the neural network combines convolutional layers, which detect local sequence motifs, with a bidirectional LSTM, which adds long-range context. This neural network assigns each base 11 independent probabilities for start-codon, coding, stop-codon and intergenic labels (Fig. 1a). A decoder then turns these probabilities into gene predictions through a dynamic programming approach.

**Fig 1.**
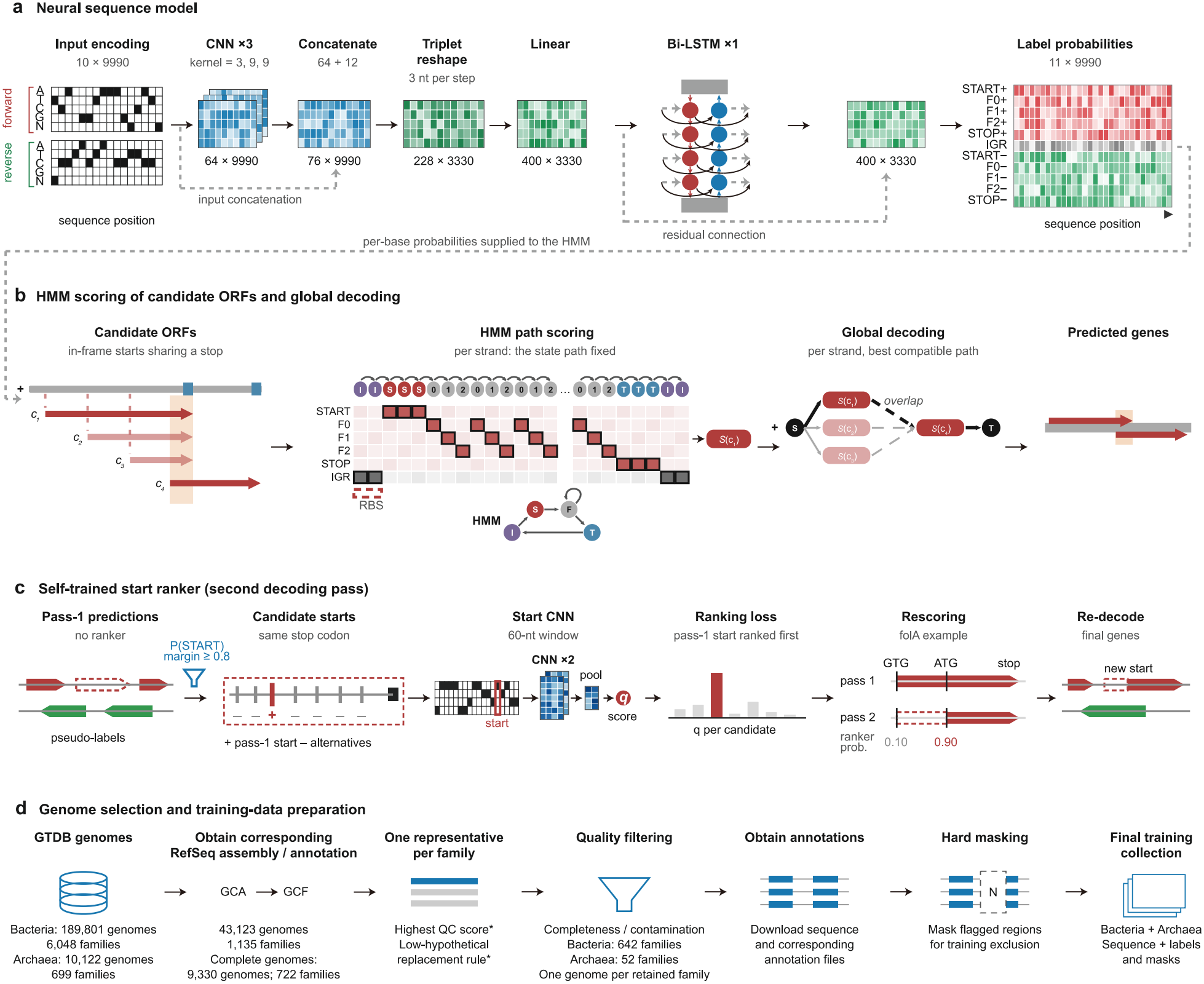
Overview of deeprok. (a) Network architecture. The input is a 9,990-nt window, one-hot encoded on both strands. Three convolutional layers and a bidirectional LSTM output a probability at each base for each of 11 labels: START, three coding frames and STOP on each strand, and intergenic. (b) Decoding. A strand HMM uses the per-base probabilities as emission probabilities to score each candidate ORF and the gap or overlap between consecutive ORFs. The highest-scoring path through the candidates gives the predicted genes, which may overlap. (c) Start refinement. A start ranker rescores the alternative in-frame start codons of each predicted gene. (d) Selection of training genomes. From the GTDB representative genomes, we kept one complete RefSeq genome per family, chosen by CheckM2 quality and the fraction of hypothetical proteins. We masked pseudogenes, non-coding RNA genes and mobile elements with N.

Two design choices allow deeprok to predict overlapping genes. First, base labels are independent: a base can be coding on both strands, or be the stop codon of one gene and the start codon of the next. Second, the decoder scores whole candidate genes rather than single bases, so neighboring genes can share bases (Fig. 1b). Each candidate ORF is scored by the log-probability of its start–coding–stop state path under the HMM, which uses the neural network’s output probabilities. To further refine start-codon assignment, deeprok adds a second-pass model. After the first-pass prediction, it trains a second, smaller neural network to rank start sites by self-training on the genome’s own confident first-pass predictions and then decodes the genome again (Fig. 1c).

We trained the first-pass network once and applied it to new genomes without retraining (Fig. 1d, Supplementary Table 1). The training set has 666 genomes: 620 bacteria and 46 archaea from 58 phyla, with one complete genome per GTDB family [15]. We also masked pseudogenes, non-coding RNAs and mobile elements during training. These segments tend to carry characteristics of genes but are not always intact protein-coding genes. We masked these segments rather than labeling them, because labeling them as either coding or intergenic sequence would give the network contradictory examples.

No benchmark genome has a training genome from the same species (Table 1). The nearest training genome of each benchmark genome is of a different genus and shares 49–62% average amino-acid identity (AAI); the only exception is *M. tuberculosis*, at 72% with a different species of the same genus. Our benchmark therefore tests how well the model generalizes to genomes distant from the training set (Supplementary Table 2).

**Table 1.** Benchmark genomes are phylogenetically distant from the training set. For each test genome, the closest GTDB (r232) taxon it shares with any training genome and that taxon’s rank; the nearest training genome by patristic distance on the GTDB tree; and the average amino-acid identity (AAI, %) between the two. AAI is the mean identity over reciprocal best BLASTP hits, using published thresholds [16]: ≥30% identity over ≥70% of the query length. Red marks the only genus-level pair. Genome with manually curated annotation; used for exact-match gene-level accuracy. Translation starts verified by N-terminal proteomics, from the GeneMarkS-2 benchmark [9]; used for start-codon accuracy.

| Test genome | Phylum | Deepest shared taxon | Nearest training genome | AAI (%) |
|---|---|---|---|---|
| *Bacteria* |  |  |  |  |
| *M. tuberculosis* H37Rv<sup>a,b</sup> | Actinomycetota | *Mycobacterium* (genus) | *Mycobacterium heraklionense* | 72 |
| *B. subtilis* 168<sup>a</sup> | Bacillota | *Bacillales* (order) | *Halobacillus* sp001592845 | 57 |
| *E. coli* K-12 MG1655<sup>a,b</sup> | Pseudomonadota | *Enterobacterales* (order) | *Kangiella geojedonensis* | 53 |
| *P. aeruginosa* PAO1<sup>a</sup> | Pseudomonadota | *Pseudomonadales* (order) | *Psychrobacter arcticus* | 51 |
| *Synechocystis* 6803<sup>b</sup> | Cyanobacteriota | *Microcystaceae* (family) | *Atelocyanobacterium thalassa* | 61 |
| *D. deserti* VCD115<sup>b</sup> | Deinococcota | *Deinococcales* (order) | *Meiothermus ruber* | 51 |
| *Archaea* |  |  |  |  |
| *H. salinarum* R1<sup>b</sup> | Halobacteriota | *Halobacteriaceae* (family) | *Halodesulfurarchaeum formicicum* | 61 |
| *N. pharaonis*<sup>b</sup> | Halobacteriota | *Halobacteriales* (order) | *Halorussus lipolyticus* | 62 |
| *A. pernix* K1<sup>b</sup> | Thermoproteota | *Acidilobaceae* (family) | *Caldisphaera lagunensis* | 49 |

### deeprok is more accurate than Prodigal and GeneMarkS-2 on curated annotations

We compared deeprok with Prodigal [8] and GeneMarkS-2 [9], two widely used ab initio gene finders for prokaryotes. We also tested the deep-learning gene finder Balrog [13], but it produced more exact-match errors than the other callers on the four curated genomes below, mainly because it placed starts less accurately (Supplementary Table 3). We thus compared only deeprok with Prodigal and GeneMarkS-2 for the rest of this study. Due to the sparsity of experimentally confirmed prokaryotic genome annotations, we used a different independent reference for each aspect of gene finding. To evaluate gene-level exact accuracy, we selected four manually curated annotations of phylogenetically diverse bacteria: *E. coli* K-12 (EcoCyc [17]), *Bacillus subtilis* 168 (SubtiWiki [18]), *Mycobacterium tuberculosis* H37Rv (Mycobrowser [19]) and *Pseudomonas aeruginosa* PAO1 (PseudoCAP [20]) (Table 2; Fig. 2a–c). Each is based on expert or community re-annotation that reconciled computational predictions with the experimental literature. We also adopted a larger set of 89,466 ORFs with peptide evidence in 54 genomes, compiled in the GeneMarkS-2 study, to evaluate gene detection, although start accuracy could not be assessed with this set (Fig. 2d). Start-codon accuracy was additionally measured on 2,925 genes whose N-termini were determined by N-terminal proteomics in four bacterial and three archaeal genomes, as compiled for the GeneMarkS-2 benchmark [9] (Methods; Fig. 3).

**Table 2.** Exact-match F1 against manually curated references. Total errors (false positives + false negatives) in parentheses. The highest F1 in each row is in bold.

| Genome | Reference CDS | deeprok | deeprok (no ranker) | Prodigal | GeneMarkS-2 |
|---|---|---|---|---|---|
| *E. coli* K-12 | 4,340 | **0.918 (705**) | 0.905 (811) | 0.898 (881) | 0.884 (995) |
| *B. subtilis* 168 | 4,240 | **0.905 (803)** | 0.896 (875) | 0.885 (978) | 0.898 (861) |
| *M. tuberculosis* H37Rv | 4,018 | **0.749 (2,004)** | 0.744 (2,027) | 0.733 (2,163) | 0.694 (2,460) |
| *P. aeruginosa* PAO1 | 5,586 | **0.896 (1,173)** | 0.896 (1,173) | 0.896 (1,175) | 0.876 (1,407) |
| Pooled | 18,184 | **0.871 (4,685)** | 0.865 (4,886) | 0.858 (5,197) | 0.843 (5,723) |

**Table 3.** Start accuracy on genes with experimentally verified starts. The method with the highest accuracy is in bold.

| Genome | Domain | Verified genes | deeprok | deeprok (no ranker) | Prodigal | GeneMark S-2 |
|---|---|---|---|---|---|---|
| *E. coli* K-12 | B | 769 | **97.9** | 97.0 | 97.7 | 96.5 |
| *M. tuberculosis* H37Rv | B | 701 | **94.7** | 93.8 | 89.0 | 91.0 |
| *D. deserti* | B | 384 | **97.7** | 95.1 | 87.4 | 96.1 |
| *Synechocystis* sp. PCC 6803 | B | 96 | **97.9** | 96.9 | 96.8 | 94.7 |
| *H. salinarum* R1 | A | 530 | 97.2 | 94.7 | 97.2 | **98.5** |
| *N. pharaonis* | A | 315 | **99.4** | 98.1 | 98.4 | 98.7 |
| *A. pernix* K1 | A | 130 | 86.2 | 73.6 | **97.7** | 96.9 |
| Bacteria (pooled) |  | 1,950 | **96.7** | 95.5 | 92.5 | 94.3 |
| Archaea (pooled) |  | 975 | 96.4 | 93.0 | 97.6 | **98.4** |
| All (pooled) |  | 2,925 | **96.6** | 94.6 | 94.2 | 95.7 |

**Fig 2.**
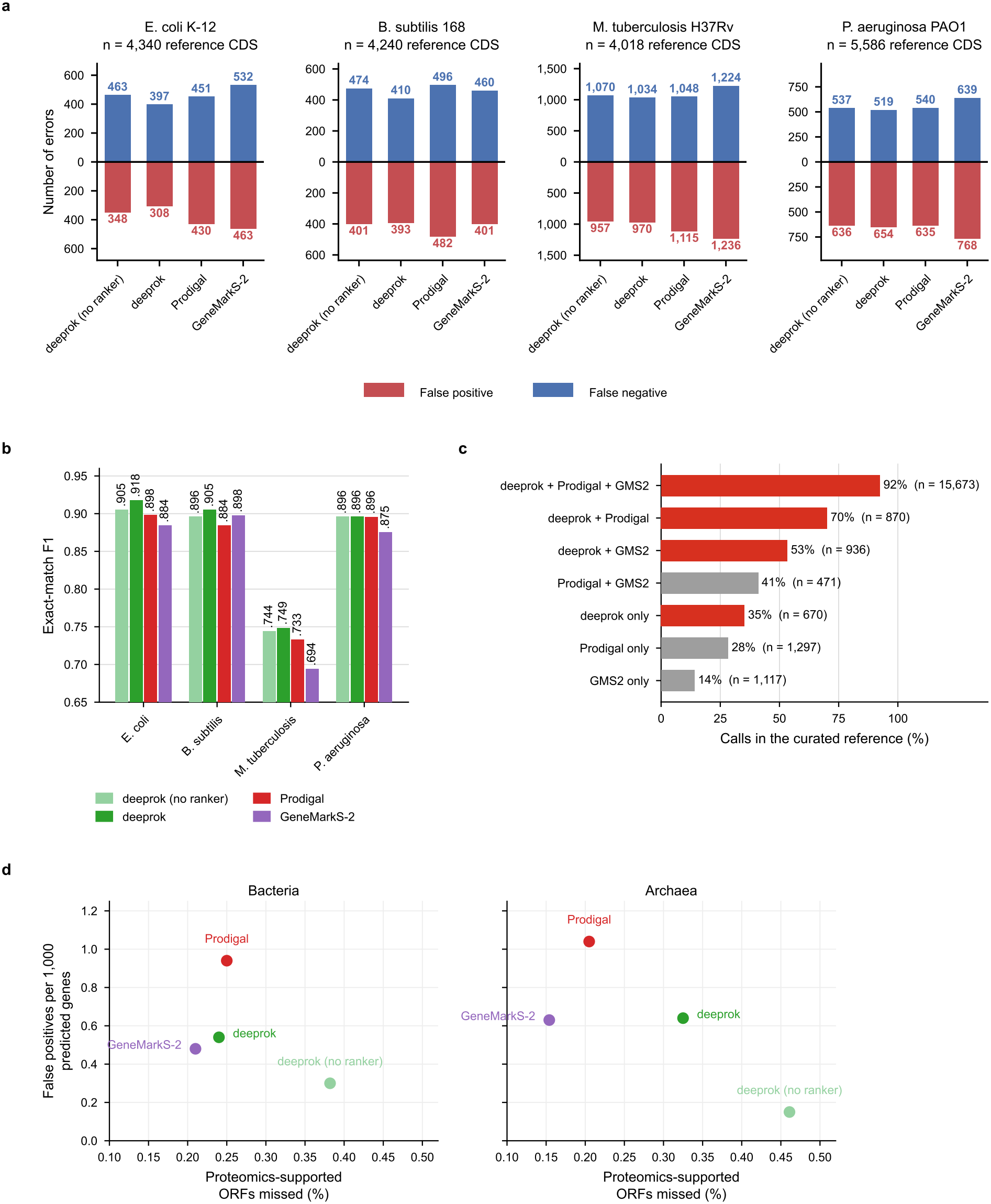
Gene-level accuracy against curated references and proteomics-supported ORFs. (a) False negatives and false positives of each caller on the four curated genomes; n is the number of reference CDS. deeprok (no ranker) is the first pass alone. (b) Exact-match F1 (both coordinates correct) on the same genomes. (c) Agreement between callers on the four curated annotations. Each bar is the set of calls made by exactly that combination of callers, and its length is the percentage of those calls in the curated reference; n is the number of calls. Red, combinations that include deeprok. (d) Proteomics-supported ORFs missed against false positives per 1,000 predicted genes, for 50 bacterial genomes (83,614 ORFs) and 4 archaeal genomes (5,852 ORFs). Lower is better on both axes.

**Fig 3.**
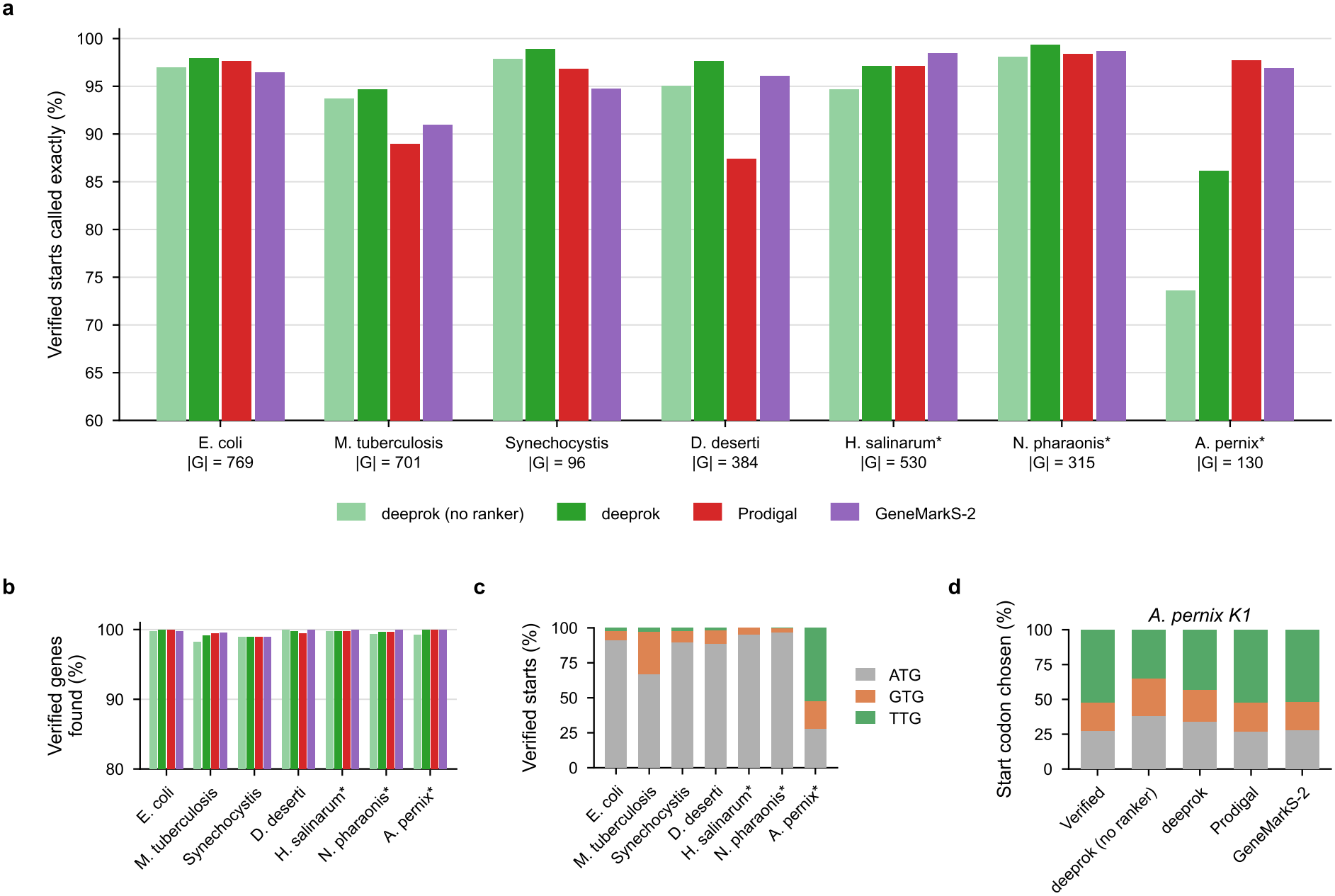
Start-codon accuracy on genomes with experimentally verified starts. (a) Start accuracy. Percentage of verified starts that each caller predicted exactly, on the seven genomes with experimentally verified starts. |G|, number of verified genes; asterisk, archaeal genome. (b) Gene recall. Percentage of verified genes that each caller found with the correct stop. (c) Start codons of the verified genes. (d) Start codons in *A. pernix* K1, in the verified set and among the starts chosen by each caller.

Pooled over all four curated genomes, deeprok had the highest exact-match accuracy (Fig. 2a, b). In each of the curated genomes, deeprok achieved similar gene recall with fewer false positives (Fig. 2a). To calculate F1, a prediction was a true positive only if its start and stop coordinates were the same as those of a reference gene. With this criterion, deeprok had the highest F1 on three of four genomes and was level with Prodigal on *P. aeruginosa. M. tuberculosis*, which has high GC content and frequent non-ATG starts [21], was the hardest genome for every caller (Fig. 2b).

A gene-by-gene comparison showed that the callers agree on most genes and differ in the reliability of their unique calls (Fig. 2c). Pooled over the four genomes, all three callers predicted 15,673 genes identically, and 92.4% of these were in the curated references. Calls unique to one caller were less often correct, but deeprok’s were the most reliable: 35.2% of its 670 unique calls were in the reference, against 28.3% for Prodigal and 14.1% for GeneMarkS-2. deeprok therefore made fewer unique calls, and a higher proportion of them were correct.

On the proteomics benchmark of 54 genomes (50 bacteria and 4 archaea), all three callers recovered more than 99.6% of the 89,466 supported ORFs, so this benchmark is close to saturation (Fig. 2d, Supplementary Table 2). deeprok missed 220 supported ORFs, against 221 for Prodigal and 185 for GeneMarkS-2, and was slightly less sensitive on the four archaeal genomes (19 missed ORFs, against 12 and 9). Thus, deeprok is more accurate than Prodigal and GeneMarkS-2 at exact gene-level prediction on curated annotations, and performs similarly to them on proteomics-supported ORFs.

### deeprok has the highest start-codon accuracy on verified N-termini

Start codons are the hardest part of prokaryotic gene prediction, and their positions in the reference gene annotation are also less certain [7]. We therefore evaluated start-codon accuracy separately, on 2,925 genes in seven genomes whose translation starts were verified by N-terminal proteomics, following the criteria used in the StartLink and GeneMarkS-2 studies [7,9]: the fraction of verified genes found with the correct stop codon that also have the correct start codon (Methods). deeprok predicted start codons more accurately than GeneMarkS-2 and Prodigal. Of the verified genes that each caller found, deeprok made only 99 start errors, against 126 for GeneMarkS-2 and 169 for Prodigal (Table 3). All three callers found 97–100% of the verified genes (Fig. 3b), so these differences reflect start selection rather than gene detection.

Most of the advantage came from the four bacterial genomes, where deeprok placed 96.7% of starts correctly, compared with 94.3% for GeneMarkS-2 and 92.5% for Prodigal. The gap was widest on *M. tuberculosis* and *Deinococcus deserti* (Table 3). In the three archaeal genomes, deeprok was less accurate than both other callers, with most errors coming from the crenarchaeon *Aeropyrum pernix*. On *A. pernix*, deeprok placed only 86.2% of starts correctly, against 97.7% for Prodigal and 96.9% for GeneMarkS-2. More than half of the verified *A. pernix* genes start with TTG (Fig. 3c), but only 43.1% of the starts deeprok predicted in this genome were TTG (Fig. 3d). deeprok therefore under-predicted TTG starts here.

The self-trained ranker improved start accuracy on every one of the seven genomes. Across all seven genomes, it reduced start errors from 156 to 99, with the largest gain on *A. pernix* (73.6% to 86.2%). The ranker partly corrects the network’s bias against TTG: the fraction of predicted start codons that are TTG increased from 27.5% to 33.5%.

Verified starts use explicit experimental evidence and exclude genes with uncertain starts, so they may give a fairer estimate of start error than the curated whole-genome references used above. Against the whole-genome curated genes, deeprok’s start-error rate appears to be 4.3% on *E. coli* and 22% on *M. tuberculosis*. On verified starts, however, it is only 2.1% and 5.3%, respectively. This discrepancy suggests that much of the apparent start error against whole-genome references may reflect uncertainty in the annotations themselves.

### deeprok’s predictions are interpretable

To see which sequence features each pass relies on, we used in silico mutagenesis (ISM) [22]: we changed each base near the curated reference start codons of the four curated genomes and measured the change in the network’s START output (without ranker) and in the ranker score. We call this change the attribution of that base (Methods). In the first pass, the network recognizes a start codon mainly by the codon itself and uses the ribosome-binding site only as supporting information (Fig. 4a). In all four genomes, the start codon had the highest attribution, especially the TG at positions +1 and +2. The Shine–Dalgarno region (7–14 nt upstream of the start codon) contributed 12–16% of the summed absolute attribution over the analysis window. The same pattern held at the 2,914 experimentally verified starts in seven genomes (9–18%).

**Fig 4.**
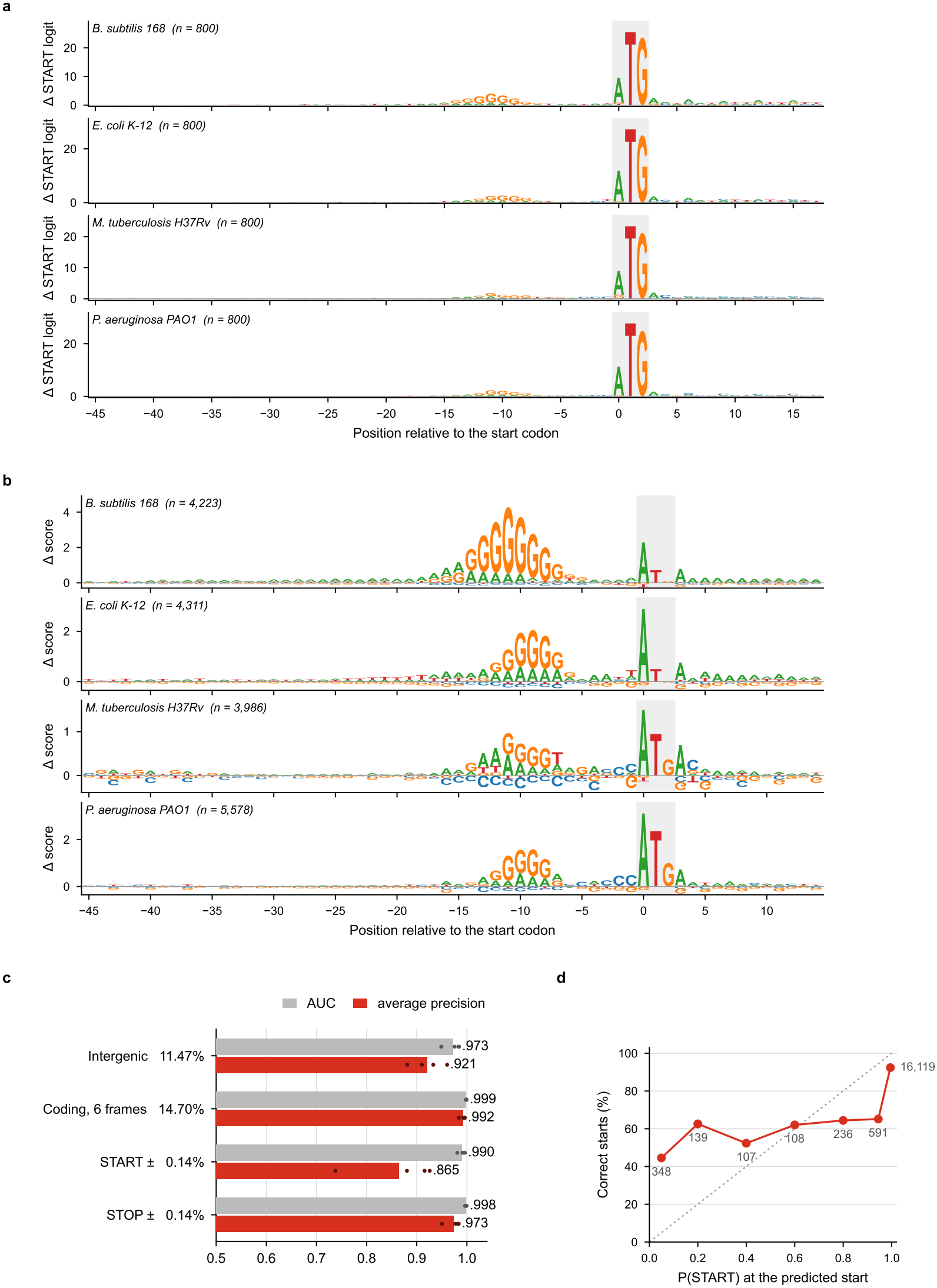
Sequence signals used by the network and the start ranker. (a) In silico mutagenesis of the network. Mean drop in the START logit when each base is replaced by each of the other three bases, around curated reference starts in the four curated genomes; n, number of starts. The start codon is shaded. (b) In silico mutagenesis of the start ranker, as in a. (c) Per-base accuracy of the network on the four curated annotations. AUC and average precision, with the 11 channels pooled into four groups; the percentage after each group is the fraction of bases that are positive for it. (d) Calibration of the start probability. Of the genes with a correct stop, the percentage with the exact start, binned by P(START) at the predicted start. The number beside each point is the number of genes in the bin; the dotted line marks perfect calibration.

On *A. pernix*, where most verified starts are TTG, the network gave TTG a mean START logit of only 1.1, consistent with its under-prediction of TTG starts in this genome.

In the second pass, the attribution of the start ranker showed a ribosome-binding-site-like pattern in each genome (Fig. 4b). In all four curated genomes, the ranker gave high attribution to a G-rich, Shine–Dalgarno-like motif approximately 7–15 nt upstream of the start codon, accounting for 29–47% of the total attribution. Interestingly, the differences between genomes were in line with known differences in translation initiation for these genomes. In *B. subtilis*, it was a long GGAGGG-like sequence with higher attribution than the start codon, consistent with the strong dependence of translation initiation in Firmicutes on Shine–Dalgarno pairing [23]. In *E. coli* and *P. aeruginosa*, the pattern was a shorter GGG sequence with lower attribution than the start codon. In *M. tuberculosis*, where a substantial fraction of mRNAs are leaderless [24,25], the motif had the lowest attribution of the four genomes.

To see how well each channel identifies its target label at single-base resolution, we computed per-base AUC and average precision against all other bases (Fig. 4c). On *E. coli* K-12, the six coding-frame outputs had an AUC of 0.998, so the network separates coding from non-coding sequence and identifies the reading frame almost perfectly. The start and stop outputs were nearly as accurate (AUC 0.992 and 0.996), even though each start or stop channel is positive at only 0.14% of bases.

Finally, we tested whether the network’s start probability, P(START), can be read as confidence that a predicted start is correct (Fig. 4d). Among genes with a correct stop, 91% had P(START) ≥ 0.99 at the predicted start, and 92.4% of those starts were exact. For the remaining 9% of calls, which scored below 0.99, start accuracy was 45–65% and did not change much with the score. Therefore, a 0.99 cutoff can still pick out uncertain starts, but lower values should not be read as true probabilities.

### Most remaining errors are misplaced starts

Most of deeprok’s remaining exact-match errors are misplaced starts (Fig. 5a). The same holds for Prodigal and GeneMarkS-2, so start placement is still a bottleneck shared by all three callers. Specifically, deeprok’s start errors were mostly short range and concentrated at non-ATG start codons. Of these errors, 68% placed the start downstream of the reference start, truncating the predicted proteins, and 45% fell within 10 codons of the true start (Fig. 5b). deeprok misplaced non-ATG starts more often than ATG starts, most often by choosing an ATG over a true GTG (Fig. 5c). Importantly, deeprok’s errors were not explained by distance to the training data (Fig. 5d). Across 12 genomes, exact-match F1 did not increase with average AAI to the nearest training genome.

**Fig 5.**
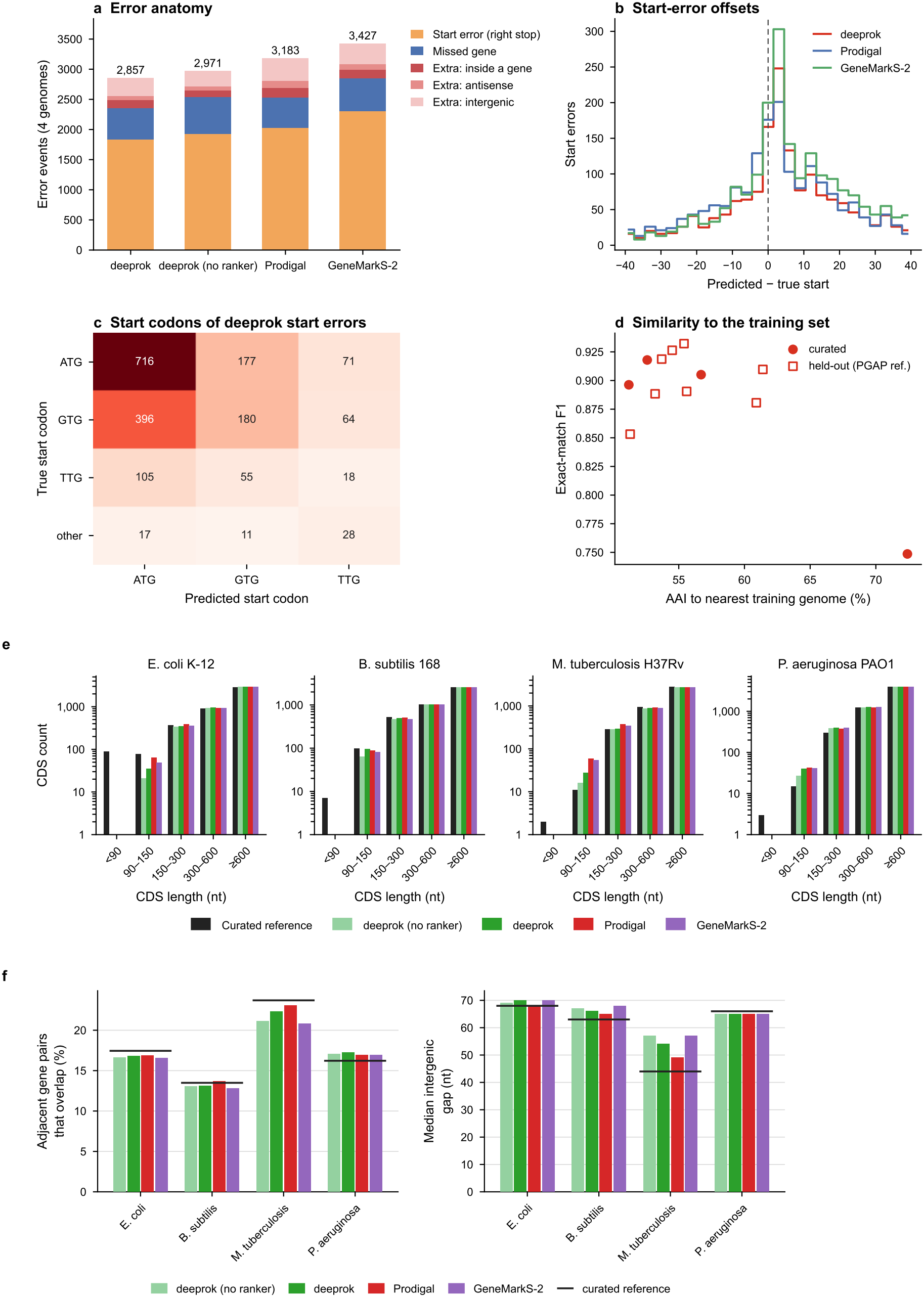
Remaining errors. (a) Error types, pooled over the four curated genomes. Extra calls are predictions with no matching stop, split by whether they overlap an annotated gene on the same strand, on the opposite strand, or no gene. (b) Start-error offsets. For genes with a correct stop but a wrong start, the distance from the true start to the predicted start, in codons (3 nt). Errors beyond ±40 codons are not shown. (c) Start codons of deeprok’s start errors. True against predicted start codon, pooled over the four curated genomes. (d) Similarity to the training set. deeprok’s exact-match F1 against the average amino-acid identity (AAI) between each benchmark genome and its nearest training genome. (e) CDS length distributions on the four curated genomes. (f) Gene-packing statistics. Percentage of adjacent gene pairs that overlap (left) and median intergenic distance (right). Black lines show the curated reference.

Besides start errors, deeprok made fewer unexplained false positives than Prodigal or GeneMarkS-2 (Supplementary Table 4). To investigate the source of false positives, we further divided extra calls by whether they lie antisense to an annotated gene, inside one on the same strand, or between genes. Across the four curated genomes, deeprok made the fewest calls antisense to a known gene (Fig. 5a). In these cases, the reverse complement of a coding sequence often contains long “shadow” ORFs, especially in GC-rich genomes, because the reverse complement rarely contains the AT-rich stop codons. Many of the other false positives, especially intergenic ones, fall on pseudogenes, which deeprok never learned to reject because we masked them during training (Supplementary Table 1). Inside an annotated gene, all callers made the same error in selenoproteins FdnG, FdoG and FdhF [26]: every caller ended these genes at the UGA codon that encodes selenocysteine. Once we set aside calls on pseudogenes, non-coding RNAs and mobile elements, deeprok had fewer unexplained false positives than either caller (281, against 394 for Prodigal and 334 for GeneMarkS-2).

We further investigated whether these unexplained calls correspond to protein-coding genes absent from the curated annotation. For each species, we searched these calls against the RefSeq proteomes of 144–150 other complete genomes of the same species (Methods). Of deeprok’s 281 unexplained false positives, 150 matched a protein annotated at full length in at least half of these genomes, compared with 177 of 394 for Prodigal and 209 of 334 for GeneMarkS-2. Most of deeprok’s matched calls were in *P. aeruginosa* (101 of 155) and *M. tuberculosis* (38 of 84), and none were in *E. coli* (0 of 6). These calls could be genes missing from the curated annotations, or systematic false positives shared by gene callers, because the other genomes were annotated with PGAP, which itself predicts genes with GeneMarkS-2+ [27].

For all three callers, missed genes were concentrated among short genes (Fig. 5e). Proteins shorter than 100 amino acids made up 1,833 of the 18,184 reference genes but accounted for 386 of deeprok’s 522 missed genes, 364 of Prodigal’s 511 and 397 of GeneMarkS-2’s 549. deeprok missed 11 more genes than Prodigal overall. The difference came from proteins of 50– 100 amino acids (185 versus 163), whereas between 100 and 300 amino acids deeprok missed fewer genes (127 versus 138). Proteins longer than 300 amino acids were almost never missed (8–9 per caller). Overall, we found no systematic bias in deeprok’s gene-packing statistics relative to the other callers and the reference (Fig. 5f).

## Discussion

We built deeprok to address persistent errors in prokaryotic gene calling by using modern deep learning techniques. A network trained once on 666 genomes labels every base, an HMM scores candidate genes from these labels, and a ranker trained on each genome’s own predictions refines the starts. On curated annotations and on experimentally verified starts, deeprok made fewer errors than Prodigal and GeneMarkS-2. On the proteomics benchmark, which measures only ORF detection, all three callers were close to saturation. deeprok needs no reference sequences or retraining, and runs on a laptop CPU in minutes (Supplementary Table 5).

Unlike Helixer and Tiberius, which use an HMM to infer one state for each base [10,11], deeprok uses the HMM only to score candidate genes, and a dynamic program then selects a set of genes that may overlap. We chose this approach rather than directly using an HMM to infer the underlying states because a single state per base cannot represent two genes that share bases in a standard model; we found that a per-base Viterbi decoding of our network output was less accurate in our tests. One cost of this design is that deeprok reports a single gene set but no principled measure of how likely each predicted gene is. An overlap-aware probabilistic model may enable more principled statistics.

A key aspect of deeprok is the two-stage approach to prediction. The start ranker in the second stage uses the same idea of self-training as GeneMarkS-2 and Prodigal but differs in what it learns. Rather than relying on a discrete set of predefined classes, deeprok’s ranker reads the sequence around each candidate start and learns which features distinguish confident first-pass starts from in-frame alternatives. Without predefined motifs, it can capture lineage-specific initiation signals (Fig. 4b), which may matter most for genomes whose signals do not fit predefined classes, and for discovering novel regulatory signatures.

Earlier deep-learning gene callers for prokaryotes, such as Balrog and GeneLM, score each candidate ORF as a whole [13,14]. Because they output only one score per ORF, the sequence features behind a call are hard to locate. In contrast, deeprok predicts labels for every base, allowing its outputs to be examined at single-base resolution. A key benefit that we showed is how interpretability approaches, such as in silico mutagenesis, explain which biological motifs the network and the ranker use to recognize a start codon. The same analysis could be applied to a predicted gene with no homologs in protein databases, to show whether the prediction depends on a start codon with an upstream ribosome-binding site, on the coding pattern inside the gene, or on neither.

deeprok has several limitations, and each points to future work. First, it is less accurate on archaea, which made up only 46 of its 666 training genomes. These capture only a fraction of known archaeal diversity and adding more archaeal training genomes may help, as may training a separate model on archaea alone. Second, most of its remaining errors are misplaced starts, concentrated at non-ATG start codons, to which the network assigns low probability (most strikingly TTG in *A. pernix*). The network is trained to label each base on its own and never directly chooses among the in-frame starts of a gene. The ranker can only partially correct this bias, as it learns from confident first-pass predictions and inherits their errors. Third, we masked pseudogenes and mobile elements during training since they look like genes but are often not intact coding sequences, so labeling them as either coding or intergenic would give the network contradictory examples. Consequently, the network never learned to reject pseudogene fragments, and genuine transposase genes were also excluded. In future releases, labeling pseudogenes as their own class rather than masking them could address both issues. Fourth, short genes remain the main source of missed genes for each caller, and deeprok does not predict genes shorter than 90 nt. Small proteins differ from typical proteins in length and in sequence [28]. Since they are short and few, they play a small role in the annotations the network learns from, and few have been confirmed by ribosome profiling or proteomics. Training a larger model on the same annotations is therefore unlikely to close this gap. Future methods may instead need to build in what is known about the biology of small proteins.

## Conclusions

We have developed deeprok, a new deep-learning gene prediction method for bacteria that requires no reference sequences, experimental data or retraining, and runs on a laptop CPU in minutes. deeprok utilizes a base-level neural network with a dynamic programming approach for ORF finding, along with a self-trained start site prediction method. We have shown that deeprok outperforms existing methods on several curated and experimentally verified annotated genomes that are highly dissimilar to our training genomes. In summary, deeprok shows that combining massive data, deep-learning techniques, and biologically informed design leads to a practical path forward to improving the fundamental task of prokaryotic gene prediction.

## Methods

### Training genomes

We selected training genomes from GTDB release 232 [15]. We considered only GTDB species representatives with a RefSeq assembly at the “Complete Genome” level and a RefSeq GFF annotation, and selected one genome per GTDB family. In each family, we chose the genome with high CheckM2 [29] completeness, low contamination and few hypothetical proteins (Supplementary Methods). This gave 642 bacterial and 52 archaeal genomes. Of these 694 genomes, we used 666 (620 bacteria and 46 archaea, from 58 phyla) as one combined training set, 14 (12 bacteria and 2 archaea) for validation and 14 (10 bacteria and 4 archaea) as held-out test genomes.

*E. coli* K-12 MG1655 has one of the best curated bacterial genome annotations. To keep this benchmark independent of training, we excluded its whole family, Enterobacteriaceae, from the training and validation sets.

Before training, we hard-masked the loci that the network cannot learn as protein-coding genes. For these loci, we replaced the sequence with N and removed the annotation. Specifically, we masked rRNA, tRNA, tmRNA and ncRNA genes, genes marked as pseudogenes, and CDS whose product name identifies a mobile element (transposase, transposon, insertion sequence, integrase or recombinase). We also removed each CDS that then contained a masked base, so that no label includes an N. We kept genes annotated as hypothetical proteins. In the training and validation genomes, masking removed 138,286 annotations. We did not mask the test genomes during inference.

### Input encoding and labels

We encode each genomic position as 10 channels. The first five channels are a one-hot encoding of the forward strand (A, C, G, T and N). The last five channels are the same encoding of the complementary base. Thus, the network reads both strands in one pass, in genomic coordinates.

The labels are a multi-hot matrix with L rows and 11 channels: channel 0 is intergenic; channels 1–3 and 4–6 are the three codon positions of coding sequence on the forward and reverse strands; channels 7 and 8 are start codons and channels 9 and 10 are stop codons, each forward then reverse.

In each CDS, the first three bases have the start-codon label and the last three bases have the stop-codon label. These bases do not have a frame label. Each other base has a label for its position in the codon (channels 1–3 or 4–6), counted from the first base of the start codon. We label reverse-strand genes on the reverse complement and then map the labels back to genomic order. Labels from different genes are added, not replaced. Thus, a base in two genes, on the same strand or on opposite strands, has the labels of both genes. The intergenic channel is set only where no other channel is set.

### Network architecture and training

The network has two main parts: convolutional layers that detect local sequence motifs, and a bidirectional LSTM that adds long-range context. Similar combinations of convolutional and recurrent layers have been used for eukaryotic gene prediction [10,11]. The network gives 11 independent outputs for each base, for both strands at once, so that a base shared by two overlapping genes can carry the labels of both. We trained it without an HMM layer and used the HMM only for decoding (see Decoding). With 1.1 million parameters, the network is small enough to annotate a bacterial genome on a laptop CPU in minutes. Supplementary Methods lists the layers and their sizes.

We divided the genomes into windows of 9,990 nt, with a step of 4,995 nt. In each training epoch, we sampled 500 windows from each genome with replacement (333,000 windows in total). Thus, each genome contributed equally to training, independent of its size.

We trained the network with AdamW (learning rate 2 × 10^−4^, weight decay 10^−5^, batch size 8) on one NVIDIA H200 GPU. We stopped training when the validation loss on the 14 validation genomes did not decrease for 15 epochs (minimum 10 epochs). We used the checkpoint with the lowest validation loss (epoch 13 of 28) for all analyses. The full run took approximately 42 h (approximately 19.5 h to the selected checkpoint). The model has 1,142,529 parameters.

#### Training loss

Earlier deep-learning gene callers were designed for eukaryotic genomes, in which coding sequence covers only a small fraction of the genome, and their losses address that imbalance [10,11]. Prokaryotic genomes pose different problems. Most bases are coding (86% in *E. coli* K-12), but gene boundaries are rare: each start-codon channel is positive at only 0.14% of bases. Neighboring genes often overlap, so one base can belong to two genes. Short genes are hard to detect [5], and with a per-base loss each gene contributes in proportion to its length, so short genes carry little weight. We designed the loss around these properties. We used a binary cross-entropy on each channel, so that a base can carry more than one label. We gave the start- and stop-codon channels a larger weight and added an F1-loss computed only on these channels. Finally, we gave the bases of short genes a larger weight, so that each gene shorter than 300 nt has the same total weight as a 300-nt gene.

We use the notation of [11]. For a window of length T, *Y*[*j, i*] ∈ 0, 1 is is the label of position j for channel, i ∈ {0, …, 10}*Z*[*j, i*] is the logit, and Ŷ [*j, i*] =*σ*( *Z*[*j, i*] the predicted probability,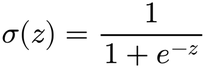where is the logistic sigmoid. The training loss combines a binary cross-entropy (BCE) with the F1-loss, whose terms we define in the paragraphs below:

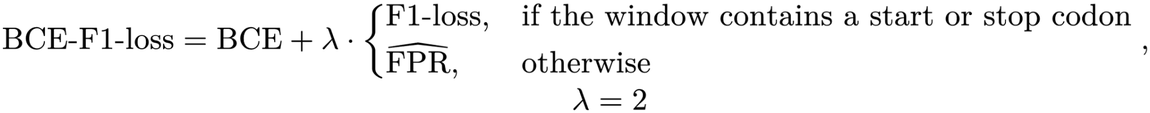

The first term is a weighted binary cross-entropy:

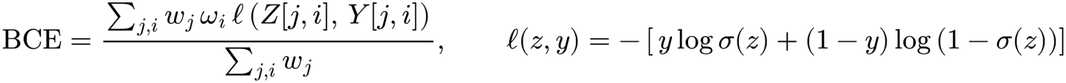

Each term has two weights: a position weight *w*_*j*_, set by the length of the gene that contains position j, and a channel weight *ω*_*i*_. In training, we calculated the BCE over all windows in a batch at once: both sums run over every position of every window.

Because start and stop codons are sparse, their errors carry little weight in an unweighted loss. Thus, we gave the four boundary channels, S = {7, 8, 9, 10}, a larger channel weight *ω*_*i*_:

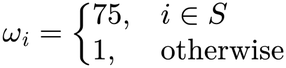

The position weight depends on the length L_g (nt) of the CDS g that contains position j. With this weight, each gene shorter than 300 nt has the same total weight as a 300-nt gene. The weight cannot be more than 5, so this applies down to 60 nt. Genes of 300 nt or longer and intergenic bases have weight 1:

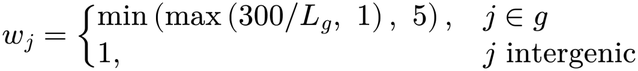

The second term is an F1-loss [11], computed only on the start-codon and stop-codon channels (the set S). We calculated it separately for each window and averaged it over the windows in a batch. For a window with at least one start-codon or stop-codon base, the term is the sum of 1 − F1 over the channels in S that have a positive base. For channel i, F1 is a soft F1 score computed from the predicted probabilities:

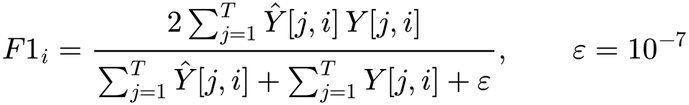

The numerator is twice the predicted probability that falls on true boundary bases. The denominator is the total predicted probability in the channel plus the number of true boundary bases. A base that is correctly predicted as not a boundary (Y[j, i] = 0 and Ŷ[j, i] close to 0) adds nothing to either sum. Therefore, the F1-loss depends only on the true boundary bases and on where the network places boundary probability. In the BCE, by contrast, boundary bases are only a small fraction of the terms, so without the channel weight ω_i their errors would have little effect. For a window without a start-codon or stop-codon base, the term is the estimated false-positive rate: the mean predicted probability, summed over the four channels in S. Training therefore decreases the predicted start-codon and stop-codon probabilities in these windows.

As in Helixer and in the pre-HMM training phase of Tiberius, we computed the loss on the network’s output probabilities and used the HMM only for decoding [10,11].

### Decoding

We used the strand HMM as a probability model for scoring candidate genes, not for inferring hidden states. Standard HMM decoding, as in Tiberius and Helixer, assigns each base a single state, either with the Viterbi algorithm or from posterior state probabilities [10,11]. A single state per base cannot represent two genes that share bases, which is common in prokaryotic genomes: at an overlap, the posterior probability of each base is divided between the two genes. In our tests, per-base Viterbi decoding of the network output was less accurate than the decoder described here, and computing posterior probabilities over whole genomes was slow. We therefore listed candidate genes and scored each by the log-probability of its own state path under the HMM. Scoring along a state path forces each candidate to follow valid codon structures and penalizes improbable transitions. A dynamic program then selects among these candidates (below).

We decoded each strand separately. Each strand uses an HMM with one intergenic state, three start-codon states, three coding states (one for each codon position) and three stop-codon states. We used the HMM notation of Tiberius [11]: Q is the set of states, *X* = *f*(*S*) ∈ [0, 1]^*T*×11^ is the network output for a sequence S of length T (Ŷ in the training loss), and X[i, q] is the output at position i for the channel of state q.

In each reading frame, each stop codon defines a group of candidates. The group contains all in-frame ATG, GTG and TTG codons after the previous in-frame stop codon. We removed candidates shorter than 90 nt or longer than the maximum gene length (125,000 nt). A candidate gene g is one start codon of a group together with the group’s stop codon. It runs from position u, the first base of the start codon, to position v, the last base of the stop codon, and defines one start–coding–stop state path Y_u, …, Y_v ∈ Q. We scored g by the log-probability of this path in the HMM:

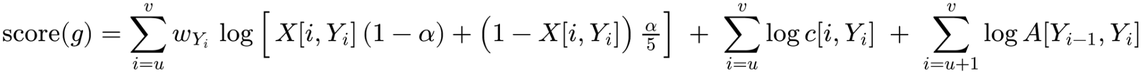

This is the log-probability of the gene’s state path in the strand HMM, with one addition: the weight w_q on the start-codon emissions. The first term is the emission from the network. As in Tiberius, each state uses only the network output for its own channel (for example, a coding state in codon position 2 uses only the frame-2 output), smoothed with α = 0.001. We multiplied this term by w_q = s for start-codon states (s = 50 in pass 1 and 10 in pass 2) and w_q = 1 otherwise. The second term is the codon factor c[i, q]. It depends on the codon that begins at position i and is 1 except at the first base of a codon: at the first start-codon state, it is the frequency of the start codon; at the first stop-codon state, it is the frequency of the stop codon; and at the first base of each internal codon, it is 0 for a stop codon and 1 otherwise, which excludes in-frame stop codons. We calculated both sets of frequencies from all 666 training genomes pooled; no benchmark genome contributes to its own priors. The third term uses the transition probabilities A[q, q′] from state q to state q′, which come from the mean gene and intergenic lengths of the training genomes.

A dynamic program over the candidates selects the chain of compatible candidates with the highest total score. The total score is the sum of the candidate scores and the connector scores between consecutive genes. A connector is either an intergenic gap or an overlap of 1 to D nt, where the maximum overlap D is 50 nt in pass 1 and 250 nt in pass 2. For an intergenic gap of n nt, the connector score is the network’s log evidence for the intergenic label summed over the gap, plus the log-probability of n under a geometric distribution whose mean is the mean intergenic length of the training genomes. For an overlap of d nt, it is the network’s log evidence that each shared base has its role in both genes (the stop codon or coding frame of the first gene, and the start codon or coding frame of the second), averaged over the two genes, plus the log-probability of d under an overlap-length prior. The prior puts most of its weight on overlaps of 1, 4 and 7 nt, the most common overlaps between prokaryotic genes [12], and decreases for longer overlaps (Supplementary Methods). Gene length enters the score only through the HMM transition scores, which correspond to a geometric length distribution; we did not add a separate gene-length prior.

Let F(g) be the best total score of a chain that ends with candidate g. The dynamic program calculates

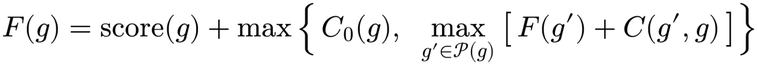

where P(g) is the set of candidates that end before g starts or overlap g by at most D nt, C(g′, g) is the connector score (intergenic gap or overlap), and C_0(g) is the score of the intergenic sequence before g when g is the first gene. The predicted gene set is the chain with the highest F(g) plus the score of the intergenic sequence after its last gene g, or no gene if the whole sequence scores higher as an intergenic sequence. The decoder returns a single highest-scoring set of genes and no posterior probabilities; we leave a fully probabilistic treatment of overlapping genes to future work.

We combined the predictions from the two strands. When two opposite-strand predictions overlapped by more than the maximum overlap, we kept the longer prediction. In pass 2, we also removed a prediction if at least 60% of its length was inside one longer prediction on the opposite strand. This step removes short ORFs on the reverse complement of a gene that is already predicted.

### Self-trained start ranker

The second pass learns the start-codon signals of the genome it annotates. It uses only the genome sequence and the pass-1 predictions. Each pass-1 gene is one training group. The start codon that pass 1 selected is the positive example. Each other in-frame ATG, GTG or TTG with the same stop codon is a negative example (median 19 per gene). We used only genes whose selected start codon has a network start-codon probability at least 0.8 higher than that of every other candidate in the group, to remove most uncertain start codons.

We represented each candidate start codon by a one-hot window of 60 nt: 45 nt upstream, the start codon and 12 nt downstream. The ranker is a small convolutional network with 7,857 parameters. It has two convolutions (24 filters, kernel size 7) with max-pooling between them, adaptive max-pooling to eight positions, dropout and two linear layers (192 → 16 → 1). For each gene, a softmax over the ranker scores of its candidates gives each candidate a probability of being the start codon. We trained the ranker with a cross-entropy loss: the negative log-probability of the positive example, averaged over genes. We held out 10% of the training genes for early stopping (settings in Supplementary Methods).

During inference, the trained ranker gives a score q to each candidate start codon in the genome. For a candidate k in stop-codon group G, we write its score as the sum of two terms,

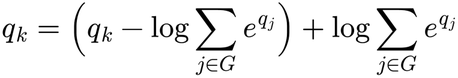

The first term, the boundary term, is the log-probability of candidate k within its group; it differs between candidates and decides which start codon is chosen. The second, the existence term, is the same for every candidate in the group; it decides whether the group is predicted as a gene or as an intergenic sequence. In pass 2, we decoded the genome again, with the score of each candidate gene g, whose start codon is candidate k of group G, replaced by

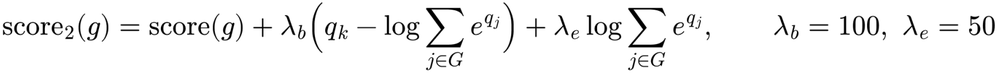

We chose λ_b and λ_e by a grid search on *E. coli* K-12 (λ_b ∈ {100, 160, 240} and λ_e ∈ {0, 50, 100, 150}), taking the pair with the highest exact-match F1, and used the same values (λ_b = 100, λ_e = 50) for all other genomes.

### Benchmark datasets and comparison callers

We used the RefSeq annotations of *E. coli* K-12 MG1655 (GCF_000005845.2) and *B. subtilis* 168 (GCF_000009045.1), the Mycobrowser release 5 annotation of *M. tuberculosis* H37Rv (NC_000962.3, without pseudogenes) and the *Pseudomonas* Genome Database annotation of *P. aeruginosa* PAO1 (NC_002516.2). The two RefSeq files are curated reference records, not PGAP predictions: every CDS in the *E. coli* file is cross-referenced to EcoCyc [30], and 95% of CDS in the *B. subtilis* file to SubtiList, the predecessor of SubtiWiki [18]. We used the experimentally verified starts compiled for the GeneMarkS-2 benchmark [9], which were determined by N-terminal protein sequencing in *Synechocystis* sp. PCC 6803 [31], *E. coli* K-12 [32], *M. tuberculosis* H37Rv [33], *D. deserti* [34], *A. pernix* K1 [35], and *H. salinarum* R1 and *N. pharaonis* [36]: seven genomes and 2,925 genes in total. We used the 89,466 ORFs with peptide evidence in 54 genomes (50 bacteria and 4 archaea) from the same benchmark [9], originally from studies at the Pacific Northwest National Laboratory [37]. We used none of these sets for training. We also measured generalization on eight bacterial test genomes against their RefSeq PGAP [27] annotation (Supplementary Table 2): all bacterial test genomes except E. coli K-12, which we scored against its curated reference, and a Mycoplasma genome that uses genetic code 4. If a gene had more than one CDS record, we used the longest as the reference.

We ran Prodigal v2.6.3 [8] in single-genome mode with default settings (-i genome.fna -f gff). We ran GeneMarkS-2 v1.0 [9] (the September 2017 GitHub release, which runs without a license key; the published v1.02 requires one), with --genome-type set to bacteria or archaea for each genome (auto for the verified-start genomes) and GFF output. We ran deeprok with the settings above. deeprok and Prodigal used the standard bacterial genetic code (translation table 11). GeneMarkS-2 identifies the genetic code from the genome.

### Evaluation metrics

For exact-match evaluation, a prediction matches a reference CDS only if it has the same sequence, strand, start coordinate and stop coordinate. We calculated precision, recall and F1 for each genome.

For each genome with verified starts, let |G| be the number of verified genes, M3 the number of verified genes for which a caller predicts a gene with the same stop codon, and M5 the number for which it predicts the same start and stop codons. Start accuracy is M5/M3, and gene recall is M3/|G|.

For proteomics-supported ORFs, we used the method of the GeneMarkS-2 benchmark [9]. A caller detects a supported ORF if one of its predictions has the same 3′ end. A prediction is a false positive if more than 30% of its length overlaps a supported ORF in a different frame. We report false positives per 1,000 predicted genes.

### Phylogenetic distance, sequence identity and model interpretation

For each benchmark genome, we calculated the patristic distance to each training genome on the GTDB release 232 maximum-likelihood trees (bac120 for bacteria and ar53 for archaea). We recorded the nearest training genome. For Fig. 5d, we compared exact-match F1 with the AAI between each benchmark genome and the nearest training genome (below).

We calculated AAI between each benchmark genome and its nearest training genome from reciprocal best BLASTP [16,38] hits between the two annotated proteomes. Hits used the thresholds in Table 1 and an e-value cutoff of 10^−5^.

For in silico mutagenesis (ISM), we changed each base around a start codon to each of the other three bases; positions are given relative to the first base of the start codon (position 0). The attribution of the observed base is the negative of the mean change in model output. For the network (Fig. 4a), we changed positions −45 to +17 at a seeded random sample of 800 curated reference start codons per curated genome, and we read the START logit of the strand of each gene from 10-kb windows. The windows were in phase with the contig coordinates, which reproduces the whole-genome outputs (Supplementary Methods). We repeated this analysis at the 2,914 verified start codons that we could score in the seven genomes with experimentally verified starts. For the ranker (Fig. 4b), we changed positions −45 to +14, the ranker’s input window, at every curated reference start codon of each curated genome (18,098 in total), and scored the changes with the ranker of that genome. The Shine–Dalgarno fraction of attribution is the sum of the mean absolute attribution over positions −14 to −7, divided by the same sum over the whole window.

### Implementation and availability

We wrote deeprok in Python with PyTorch [39]. It runs on a CPU, and a GPU is optional. It is available as a conda package with fixed dependency versions, and users can also install it with pip. For a genome FASTA file, it writes a GFF3 annotation and FASTA files of proteins and CDS (translation table 11). We tested deeprok 0.1.0 with Python 3.12 and PyTorch 2.14.1, and trained the network under Python 3.13.13. For the cross-strain analysis, we ran BLAST+ 2.17.0 [38]. We obtained CheckM2 completeness and contamination scores from the GTDB release 232 metadata [15].

We have made the source code, trained model and decoding priors available at https://github.com/susanxu3/deeprok (MIT license). The bundled model is the exact checkpoint used for all analyses (sha256 7081f5ad1b15…). Accessions of the 694 training, validation and test genomes are provided in Supplementary Table 1. Runtime and peak memory are provided in Supplementary Table 5.

## Declarations

### Authors’ contributions

Q.X. designed the method with input from H.L. and J.S. Q.X. wrote the software, performed the experiments and analyses, and wrote the manuscript. H.L. and J.S. supervised the project and revised the manuscript. All authors read and approved the final manuscript.

## Supplementary Information

### Supplementary Tables

The tables are provided as one Excel file, with one sheet per table and a README sheet with the legends.

**Supplementary Table 1. Training, validation and test genomes**. One row per genome (694 rows). Columns: RefSeq and GTDB accessions; domain; set (training, 666; validation, 14; test, 14); GTDB taxonomy from phylum to species; genome size and GC; CheckM2 completeness and contamination; number of proteins and percentage of hypothetical proteins in the RefSeq annotation; and the number of annotations masked for training (138,286 in total; test genomes were not masked). Taxonomy, size, GC and CheckM2 values are from GTDB release 232.

**Supplementary Table 2. Benchmark genomes**. One row per genome in the three benchmarks: four curated genomes, seven genomes with experimentally verified starts and 54 genomes with proteomics-supported ORFs. It also includes the eight held-out GTDB test genomes. Columns: accession; reference annotation; number of reference genes, verified starts or supported ORFs; nearest training genome on the GTDB tree and its average amino-acid identity (AAI) to the benchmark genome (data for Fig. 5d). For the 54 proteomics genomes, the table instead gives each caller’s missed ORFs and false positives (predictions with more than 30% of their length overlapping a supported ORF in another frame).

**Supplementary Table 3. Comparison with Balrog on the four curated genomes**. Exact-match and stop-level accuracy of the four callers, per genome in the Excel file and pooled over 18,184 reference genes below. Start accuracy is the fraction of genes with a correct stop that also have the correct start. The per-genome version of this table also records which genomes are in Balrog’s training or reference data: *E. coli* K-12 is in Balrog’s test set; *B. subtilis* 168 and *M. tuberculosis* H37Rv are in its protein reference database; strains of *B. subtilis, M. tuberculosis* and *P. aeruginosa* are in its training data.

**Supplementary Table 4. Classification of false-positive calls on the four curated genomes**. One row per caller and genome, plus pooled rows. Each false positive (a prediction without an exact match) is classed as correct stop but wrong start, in a reference reading frame with a different stop, in another frame on the same strand, antisense, or intergenic. Calls outside every reference frame are then flagged if they overlap an annotated pseudogene, non-coding RNA or mobile element, and the rest are unexplained (deeprok 281, Prodigal 394, GeneMarkS-2 334, pooled).

**Supplementary Table 5. Runtime and peak memory on the four curated genomes**. Wall time of each stage of the deeprok pipeline, and wall time and peak resident memory of deeprok, Prodigal 2.6.3 and GeneMarkS-2, on CPU. Each time is the median of three runs with 8 threads on an exclusive cluster node (Intel Xeon Silver 4316, 2.30 GHz), and peak memory is the maximum over runs and, for deeprok, over its stages. The network forward pass uses the 49,995-nt tiles of the released pipeline. The last column is the mean of the four genomes. The Excel sheet for Supplementary Table 5 gives the totals for eight further genomes of 1.2–9.1 Mb. On a laptop CPU (Apple M3 Pro, 4 threads), deeprok annotated *E. coli* K-12 in 6.4 min with 1.6 GB peak memory.

## Supplementary Methods

### S1. Selection of training genomes

We started from the GTDB release 232 species representatives and kept those with a RefSeq assembly at the “Complete Genome” level and a RefSeq GFF annotation. In each GTDB family we kept one genome. We scored each candidate as CheckM2 completeness minus 5 × CheckM2 contamination and first chose the genome with the highest score. We then switched to the family’s genome with the lowest percentage of hypothetical proteins if its percentage was at least 5 points lower, its score was at most 5 points lower, its completeness was at least 90% and its contamination was at most 5%. This gave 642 bacterial and 52 archaeal genomes. We split them into 666 training genomes (620 bacteria and 46 archaea), 14 validation genomes (12 bacteria and 2 archaea) and 14 test genomes (10 bacteria and 4 archaea). To keep the *E. coli* K-12 benchmark independent, we excluded the family Enterobacteriaceae from the training and validation sets. Supplementary Table 1 lists every genome.

### S2. Masking of training annotations

Before training, we replaced the sequence of the following loci with N and removed their annotation: rRNA, tRNA, tmRNA and ncRNA genes; genes marked as pseudogenes; and CDS whose product name contains transposase, transposon, insertion sequence, integrase or recombinase. We then removed every CDS that contained a masked base, so that no label covers an N. We kept CDS annotated as hypothetical proteins. Masking removed 138,286 annotations from the training and validation genomes. We did not mask the benchmark genomes.

### S3. Network architecture

The input is a window of L bases encoded as 10 channels (one-hot A, C, G, T, N for the forward strand and for its complement). The network outputs 11 logits per base. Layers, in order:

| **Layer** | **Details** | **Output shape** |
|---|---|---|
| Initial convolution | Conv1d, 10 → 64 channels, kernel 3, ReLU | L × 64 |
| Convolution block (×2) | LayerNorm, Conv1d 64 → 64, kernel 9, ReLU | L × 64 |
| Input skip | Concatenate the 10 input channels | L × 74 |
| Codon packing | Reshape 3 consecutive positions into one step | L/3 × 222 |
| Projection | Linear, 222 → 400 | L/3 × 400 |
| Bidirectional LSTM | 200 units per direction, residual connection, LayerNorm | L/3 × 400 |
| Output | Linear, 400 → 33, reshaped back to bases | L × 11 |
| Activation | Logistic sigmoid on each channel | L × 11 |

The network has 1,142,529 trainable parameters, 84% of them in the LSTM. Packing three positions into one LSTM step means each step sees exactly one codon in each frame. We used no dropout.

### S4. Training

| Setting | Value |
|---|---|
| Window, step | 9,990 nt, 4,995 nt |
| Windows per genome per epoch | 500, sampled with replacement (333,000 per epoch) |
| Optimizer | AdamW, learning rate 2 × 10<sup>−4</sup>, weight decay 10<sup>−5</sup> |
| Batch size | 8 windows |
| Loss | Weighted binary cross-entropy + F1-loss on start and stop channels |
| Channel weight, start and stop channels | 75 |
| F1-loss weight | 2 |
| Gene-length weight | min(max(300 / L_g, 1), 5) for bases of a CDS of length L_g nt; 1 for intergenic bases |
| Early stopping | Validation loss on 14 genomes; patience 15 epochs; minimum 10 epochs |
| Selected checkpoint | Epoch 13 of 28 |
| Hardware, time | One NVIDIA H200 GPU; about 42 h in total, about 19.5 h to the selected checkpoint |
| Random seed | 42 |

### S5. Decoding

At inference, deeprok runs the network on 49,995-nt tiles with half-tile overlap and decodes each strand separately. The start- and stop-codon frequencies and the mean gene and intergenic lengths for the HMM transitions were estimated once from the 666 training genomes pooled. The total probability of an overlap (15.6%) is the fraction of neighboring same-strand gene pairs that overlap in three enterobacterial genomes (E. coli O157:H7 Sakai, E. coli UMN026 and Salmonella LT2), none of which is in the training set, and its split among overlap lengths (below) was set by hand. All of these priors are the same for every genome.

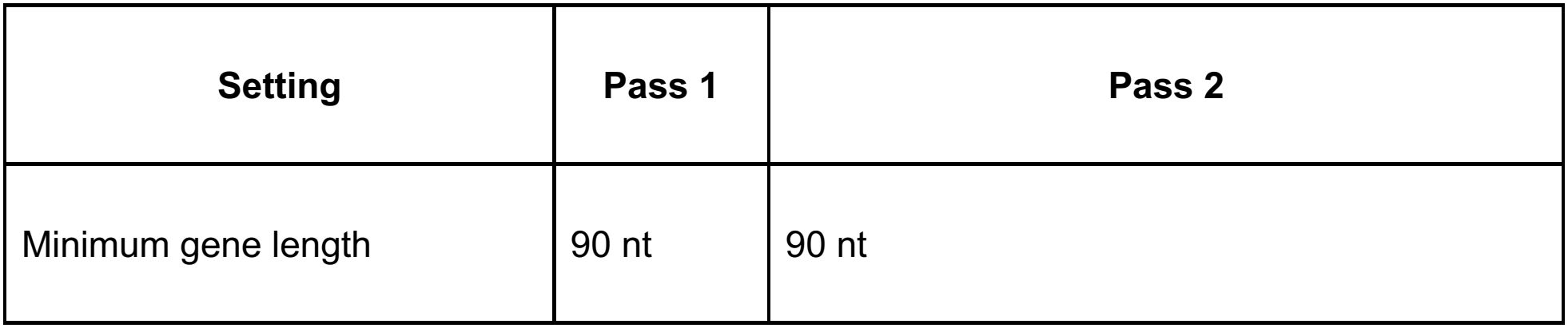
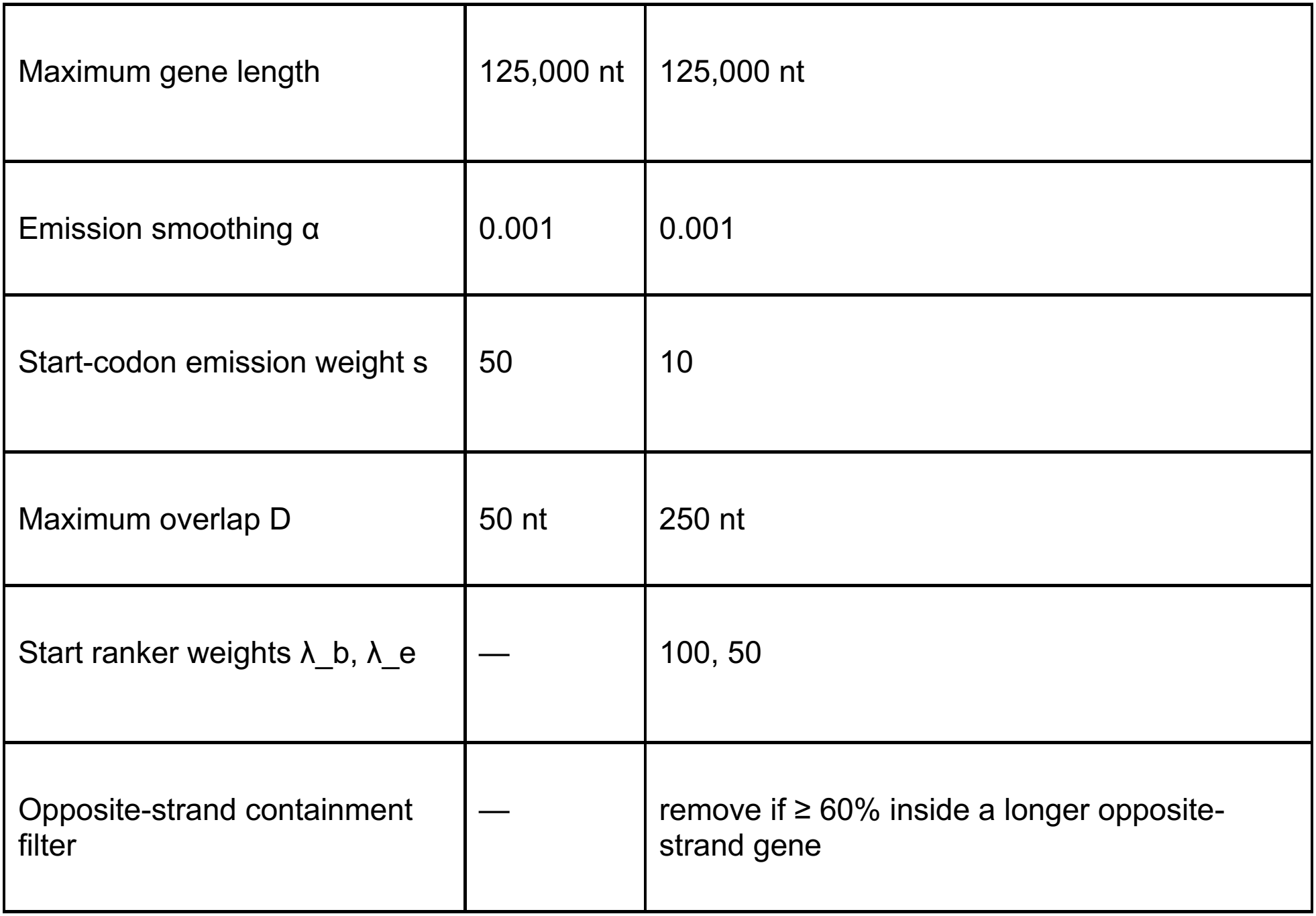

| Setting | Pass 1 | Pass 2 |
|---|---|---|
| Minimum gene length | 90 nt | 90 nt |
| Maximum gene length | 125,000 nt | 125,000 nt |
| Emission smoothing α | 0.001 | 0.001 |
| Start-codon emission weight s | 50 | 10 |
| Maximum overlap D | 50 nt | 250 nt |
| Start ranker weights λ_b, λ_e | — | 100, 50 |
| Opposite-strand containment filter | — | remove if ≥ 60% inside a longer opposite-strand gene |

Connector score. Between two consecutive genes on a strand, the decoder adds the score of the sequence between them. An intergenic gap scores the intergenic emissions over the gap plus a geometric term for its length. An overlap of 1 to D nt scores the network’s evidence at the shared bases plus the log-probability of that overlap length under the overlap-length prior. The prior splits the overlap probability 30%, 55% and 15% among overlaps of 1, 4 and 7 nt, interpolates between them on the log scale, and beyond 7 nt decreases by 1 nat per 10 nt, up to the maximum overlap D.

### S6. Start ranker

For each genome, the ranker is trained on the pass-1 genes whose selected start has a network start probability at least 0.8 higher than every other in-frame candidate with the same stop codon. Candidates are in-frame ATG, GTG and TTG codons that give an ORF of at least 30 nt, including the stop codon. Each candidate is represented by a one-hot window of 60 nt (45 nt upstream, the start codon and 12 nt downstream). The ranker has two convolutions (24 filters, kernel 7) with max-pooling between them, adaptive max-pooling to 8 positions, dropout (0.3) and two linear layers (192 → 16 → 1), with 7,857 parameters. We trained it with a listwise softmax cross-entropy over the candidates of each gene, using AdamW (learning rate 3 × 10^−3^, weight decay 10^−3^, batch 64 genes) for up to 40 epochs, with early stopping (patience 6) on 10% of genes held out. Pass 2 adds each candidate’s centered ranker score, weighted by λ_b, and the log-sum of the scores in its group, weighted by λ_e. We chose λ_b and λ_e by a grid search on *E. coli* K-12. The ranker is skipped if fewer than 100 genes pass the confidence filter.

### S7. Cross-strain analysis of unexplained false positives

For each curated genome, we downloaded the RefSeq protein sets of 144–150 other complete genomes of the same species. For each species, we took the complete RefSeq genomes, removed the reference strain and its laboratory derivatives (for example, K-12 strains for E. coli and H37Rv and H37Ra re-sequencings for M. tuberculosis), and kept the 150 with the highest CheckM completeness (ties broken by lower contamination); 144 remained for M. tuberculosis. We searched each caller’s unexplained false positives against them with blastp (e-value ≤ 10^−5^). We counted a call as matching a full-length protein in a genome if the best hit had at least 95% identity and the alignment covered at least 90% of both the query and the subject. A call was supported if it matched a full-length protein in at least half of the genomes.

### S8. Classification of false-positive calls

An extra call is a prediction whose stop codon does not match any reference gene. We classified each extra call by its overlap with reference genes: antisense (overlaps a gene on the opposite strand), same-strand (lies inside a gene on the same strand) or intergenic (overlaps no gene), where an overlap of at least 1 nt counts. Separately, we flagged calls that overlap an annotated pseudogene, non-coding RNA or mobile element. The calls left after removing these flags are the unexplained false positives in Supplementary Table 4.

### S9. Per-base accuracy and calibration

For Fig. 4c, we computed the area under the ROC curve and the average precision of each output channel on *E. coli* K-12, treating each base as positive for a channel if its label is set, and pooled the 11 channels into four groups (coding frames, start, stop, intergenic). For Fig. 4d, we took the genes whose stop was predicted correctly, binned them by P(START) at the predicted start, and computed the percentage of genes in each bin whose start was also correct.

### S10. Software versions and commands

- deeprok, deeprok -i genome.fna -o genome.gff with default settings, translation table 11.
- Prodigal v2.6.3, single-genome mode, prodigal -i genome.fna -f gff.
- GeneMarkS-2 v1.0 (September 2017 GitHub release, which runs without a license key), --genome-type bacteria or archaea (auto for the verified-start genomes), GFF output.
- Balrog 0.3.2, run with the official Colab notebook and its default settings (maximum overlap 60 nt, minimum ORF 60 nt, protein k-mer and MMseqs2 filters on, translation table 11).

## References

1. Salzberg SL. Next-generation genome annotation: we still struggle to get it right. Genome Biol. 2019;20:92. 10.1186/s13059-019-1715-2

2. Land M, Hauser L, Jun S-R, Nookaew I, Leuze MR, Ahn T-H, et al. Insights from 20 years of bacterial genome sequencing. Funct Integr Genomics. 2015;15:141–61. 10.1007/s10142-015-0433-4

3. Jundzill M, Hölzer M, Mangul S, Marquet M, Ehricht R, Lohde M, et al. Large-scale benchmarking of prokaryotic annotation tools across thousands of species. Genome Biol. 2026;27:284. 10.1186/s13059-026-04262-0

4. Dimonaco NJ, Aubrey W, Kenobi K, Clare A, Creevey CJ. No one tool to rule them all: prokaryotic gene prediction tool annotations are highly dependent on the organism of study. Bioinformatics. 2022;38:1198–207. 10.1093/bioinformatics/btab827

5. Nielsen P, Krogh A. Large-scale prokaryotic gene prediction and comparison to genome annotation. Bioinformatics. 2005;21:4322–9. 10.1093/bioinformatics/bti701

6. Gray T, Storz G, Papenfort K. Small Proteins; Big Questions. J Bacteriol. 2022;204:e0034121. 10.1128/JB.00341-21

7. Gemayel K, Lomsadze A, Borodovsky M. StartLink and StartLink+: Prediction of Gene Starts in Prokaryotic Genomes. Front Bioinform. 2021;1:704157. 10.3389/fbinf.2021.704157

8. Hyatt D, Chen G-L, Locascio PF, Land ML, Larimer FW, Hauser LJ. Prodigal: prokaryotic gene recognition and translation initiation site identification. BMC Bioinformatics. 2010;11:119. 10.1186/1471-2105-11-119

9. Lomsadze A, Gemayel K, Tang S, Borodovsky M. Modeling leaderless transcription and atypical genes results in more accurate gene prediction in prokaryotes. Genome Res. 2018;28:1079–89. 10.1101/gr.230615.117

10. Holst F, Bolger AM, Kindel F, Günther C, Maß J, Triesch S, et al. Helixer: ab initio prediction of primary eukaryotic gene models combining deep learning and a hidden Markov model. Nat Methods. 2026;23:732–9. 10.1038/s41592-025-02939-1

11. Gabriel L, Becker F, Hoff KJ, Stanke M. Tiberius: end-to-end deep learning with an HMM for gene prediction. Bioinformatics. 2024;40:btae685. 10.1093/bioinformatics/btae685

12. Johnson ZI, Chisholm SW. Properties of overlapping genes are conserved across microbial genomes. Genome Res. 2004;14:2268–72. 10.1101/gr.2433104

13. Sommer MJ, Salzberg SL. Balrog: A universal protein model for prokaryotic gene prediction. PLOS Comput Biol. 2021;17:e1008727. 10.1371/journal.pcbi.1008727

14. Akotenou G, El Allali A. Genomic language models (gLMs) decode bacterial genomes for improved gene prediction and translation initiation site identification. Brief Bioinform. 2025;26:bbaf311. 10.1093/bib/bbaf311

15. Parks DH, Chuvochina M, Rinke C, Mussig AJ, Chaumeil P-A, Hugenholtz P. GTDB: an ongoing census of bacterial and archaeal diversity through a phylogenetically consistent, rank normalized and complete genome-based taxonomy. Nucleic Acids Res. 2022;50:D785–94. 10.1093/nar/gkab776

16. Konstantinidis KT, Tiedje JM. Towards a genome-based taxonomy for prokaryotes. J Bacteriol. 2005;187:6258–64. 10.1128/JB.187.18.6258-6264.2005

17. Keseler IM, Gama-Castro S, Mackie A, Billington R, Bonavides-Martínez C, Caspi R, et al. The EcoCyc Database in 2021. Front Microbiol. 2021;12:711077. 10.3389/fmicb.2021.711077

18. Pedreira T, Elfmann C, Stülke J. The current state of SubtiWiki, the database for the model organism Bacillus subtilis. Nucleic Acids Res. 2022;50:D875–82. 10.1093/nar/gkab943

19. Kapopoulou A, Lew JM, Cole ST. The MycoBrowser portal: a comprehensive and manually annotated resource for mycobacterial genomes. Tuberculosis. 2011;91:8–13. 10.1016/j.tube.2010.09.006

20. Winsor GL, Griffiths EJ, Lo R, Dhillon BK, Shay JA, Brinkman FSL. Enhanced annotations and features for comparing thousands of Pseudomonas genomes in the Pseudomonas genome database. Nucleic Acids Res. 2016;44:D646–53. 10.1093/nar/gkv1227

21. Cole ST, Brosch R, Parkhill J, Garnier T, Churcher C, Harris D, et al. Deciphering the biology of Mycobacterium tuberculosis from the complete genome sequence. Nature. 1998;393:537–44. 10.1038/31159

22. Zhou J, Troyanskaya OG. Predicting effects of noncoding variants with deep learning–based sequence model. Nat Methods. 2015;12:931–4. 10.1038/nmeth.3547

23. Vellanoweth RL, Rabinowitz JC. The influence of ribosome-binding-site elements on translational efficiency in Bacillus subtilis and Escherichia coli in vivo. Mol Microbiol. 1992;6:1105–14. 10.1111/j.1365-2958.1992.tb01548.x

24. Shell SS, Wang J, Lapierre P, Mir M, Chase MR, Pyle MM, et al. Leaderless Transcripts and Small Proteins Are Common Features of the Mycobacterial Translational Landscape. PLoS Genet. 2015;11:e1005641. 10.1371/journal.pgen.1005641

25. Cortes T, Schubert OT, Rose G, Arnvig KB, Comas I, Aebersold R, et al. Genome-wide Mapping of Transcriptional Start Sites Defines an Extensive Leaderless Transcriptome in Mycobacterium tuberculosis. Cell Rep. 2013;5:1121–31. 10.1016/j.celrep.2013.10.031

26. Böck A, Forchhammer K, Heider J, Leinfelder W, Sawers G, Veprek B, et al. Selenocysteine: the 21st amino acid. Mol Microbiol. 1991;5:515–20. 10.1111/j.1365-2958.1991.tb00722.x

27. Tatusova T, DiCuccio M, Badretdin A, Chetvernin V, Nawrocki EP, Zaslavsky L, et al. NCBI prokaryotic genome annotation pipeline. Nucleic Acids Res. 2016;44:6614–24. 10.1093/nar/gkw569

28. Storz G, Wolf YI, Ramamurthi KS. Small proteins can no longer be ignored. Annu Rev Biochem. 2014;83:753–77. 10.1146/annurev-biochem-070611-102400

29. Chklovski A, Parks DH, Woodcroft BJ, Tyson GW. CheckM2: a rapid, scalable and accurate tool for assessing microbial genome quality using machine learning. Nat Methods. 2023;20:1203–12. 10.1038/s41592-023-01940-w

30. Karp PD, Paley S, Caspi R, Kothari A, Krummenacker M, Midford PE, et al. The EcoCyc database (2025). EcoSal Plus. 2025;13:eesp-0019-2024. 10.1128/ecosalplus.esp-0019-2024

31. Choi J-S, Kim D-S, Lee J, Kim S-J, Kim S-I, Kim YH, et al. Proteome Analysis of Light-induced Proteins in Synechocystis sp. PCC 6803: Identification of Proteins Separated by 2D-PAGE Using N-terminal Sequencing and MALDI-TOF MS. Mol Cells. 2000;10:705–11. 10.1007/s10059-000-0705-1

32. Zhou J, Rudd KE. EcoGene 3.0. Nucleic Acids Res. 2013;41:D613–24. 10.1093/nar/gks1235

33. Lew JM, Kapopoulou A, Jones LM, Cole ST. TubercuList—10 years after. Tuberculosis. 2011;91:1–7. 10.1016/j.tube.2010.09.008

34. de Groot A, Roche D, Fernandez B, Ludanyi M, Cruveiller S, Pignol D, et al. RNA Sequencing and Proteogenomics Reveal the Importance of Leaderless mRNAs in the Radiation-Tolerant Bacterium Deinococcus deserti. Genome Biol Evol. 2014;6:932–48. 10.1093/gbe/evu069

35. Yamazaki S, Yamazaki J, Nishijima K, Otsuka R, Mise M, Ishikawa H, et al. Proteome Analysis of an Aerobic Hyperthermophilic Crenarchaeon, Aeropyrum pernix K1. Mol Cell Proteomics. 2006;5:811–23. 10.1074/mcp.M500312-MCP200

36. Aivaliotis M, Gevaert K, Falb M, Tebbe A, Konstantinidis K, Bisle B, et al. Large-scale identification of N-terminal peptides in the halophilic archaea Halobacterium salinarum and Natronomonas pharaonis. J Proteome Res. 2007;6:2195–204. 10.1021/pr0700347

37. Venter E, Smith RD, Payne SH. Proteogenomic Analysis of Bacteria and Archaea: A 46 Organism Case Study. PLOS ONE. 2011;6:e27587. 10.1371/journal.pone.0027587

38. Camacho C, Coulouris G, Avagyan V, Ma N, Papadopoulos J, Bealer K, et al. BLAST+: architecture and applications. BMC Bioinformatics. 2009;10:421. 10.1186/1471-2105-10-421

39. Paszke A, Gross S, Massa F, Lerer A, Bradbury J, Chanan G, et al. PyTorch: An Imperative Style, High-Performance Deep Learning Library. arXiv. 2019. 10.48550/arXiv.1912.01703

